# Lateral Flow Assays Biotesting by Utilizing Plasmonic Nanoparticles Made of Inexpensive Metals – Replacing Colloidal Gold

**DOI:** 10.1101/2024.01.08.574723

**Authors:** Veronica A. Bahamondes Lorca, Oscar Ávalos-Ovando, Christoph Sikeler, Heini Ijäs, Eva Yazmin Santiago, Eli Skelton, Yong Wang, Ruiqi Yang, Katherine Leslee A. Cimatu, Olga Baturina, Zhewei Wang, Jundong Liu, Joseph M. Slocik, Shiyong Wu, Dongling Ma, Andrei I. Pastukhov, Andrei V. Kabashin, Martin E. Kordesch, Alexander O. Govorov

**Affiliations:** Edison Biotechnology Institute, Ohio University, Athens, Ohio 45701, United States; Departamento de Tecnología médica, Facultad de Medicina, Universidad de Chile, Santiago, Chile; Department of Physics and Astronomy, Ohio University, Athens, Ohio 45701, United States; Nanoscale and Quantum Phenomena Institute, Ohio University, Athens, Ohio 45701, United States; Faculty of Physics and Center for NanoScience (CeNS), Ludwig Maximilians University, 80539 Munich, Germany; Department of Chemistry and Biochemistry, Ohio University, Athens, Ohio 45701, United States; Institut National de la Recherche Scientifique,Varennes, Québec J3X 1P7, Canada; Chemistry Division, United States Naval Research Laboratory, Washington DC 20375, United States; School of Electrical Engineering and Computer Science, Ohio University, Athens, Ohio 45701, United States; Soft Matter Materials Branch, Materials and Manufacturing Directorate, Air Force Research Laboratory, Wright Patterson Air Force Base, Ohio 45433-7750, United States; Laboratory LP3, Campus de Luminy, Aix-Marseille University, CNRS, 13288 Marseille, France

**Keywords:** optical activity, nanoparticles, lateral flow assay, biomarkers

## Abstract

Nanoparticles (NPs) can be conjugated with diverse biomolecules and employed in biosensing to detect target analytes in biological samples. This proven concept was primarily used during the COVID-19 pandemic with gold NPs-based lateral flow assays (LFAs). Considering the gold price and its worldwide depletion, here we show that novel plasmonic nanoparticles (NPs) based on inexpensive metals, titanium nitride (TiN) and copper covered with a gold shell (Cu@Au), perform comparable or even better than gold nanoparticles. After conjugation, these novel nanoparticles provided high figures of merit for LFA testing, such as high signals and specificity and robust naked-eye signal recognition. To the best of our knowledge, our study represents the 1st application of laser-ablation-fabricated nanoparticles (TiN) in the LFA and dot-blot biotesting. Since the main cost of the Au NPs in commercial testing kits is in the colloidal synthesis, our development with TiN is very exciting, offering potentially very inexpensive plasmonic nanomaterials for various bio-testing applications. Moreover, our machine learning study showed that the bio-detection with TiN is more accurate than that with Au.

---

The recent worldwide pandemic taught us that we were not fully prepared to undergo immediate massive and cheap preventive testing, which simultaneously needed to be reliable and easy-to-use. Nanotechnology and nanomaterials represent a still fully undiscovered frontier of novel potential uses applicable to advanced medical treatments, both for diagnostics and therapy. At the heart of all this technology are plasmonic nanoparticles (NPs), generally gold NPs (Au NPs),^1^ since diverse biomolecules can bind to the Au NP’s surface, serving as a vehicle for antibodies^2^ and allowing nanophotonic biosensing.^3,4^

Additional to the Au NPs-based COVID-19 tests market,^5–7^ colorimetric testing systems for other biomarkers and biomolecules are also commercialized in the so-called lateral-flow assays (LFAs).^8–10^ LFAs are inexpensive, quick and a reliable way of analyzing diverse biomarkers such as for pregnancy, diabetes, and strokes, among many others (see Fig. 1A). Hence, LFAs should be ideal for large-scale testing to face eventual new pandemics or for everyday illnesses testing. As such, recent studies have focused on improving Au-based-LFAs. For example, SARS-CoV-2-conjugated fluorescent gold nanorods were recently shown to achieve 95% sensitivity and 100% specificity when reading with a benchtop fluorescence scanner.^11^ However, gold is an expensive material and its production is plateauing,^12^ where the cost of Au NPs prepared by colloidal chemistry is estimated to be ∼10^5^ higher than the bulk material,^13^ and with some predictions anticipating its possible exhaustion by 2050.^14^ Additionally, the demand of metallic NPs market is soon expected to double, from ∼USD 2.4 billion in 2021 up to ∼USD 4.2 billion in 2030.^15^ So, in a market dominated with noble metal NPs,^8^ alternatives are needed, better yet if they are inexpensive and abundant for having more affordable technologies. Then, custom-made optically-active plasmonic NPs can provide a suitable alternative for current over-the-counter testing (see Fig. 1E), especially in the field of nanomedicine.^16^

**Figure 1.**
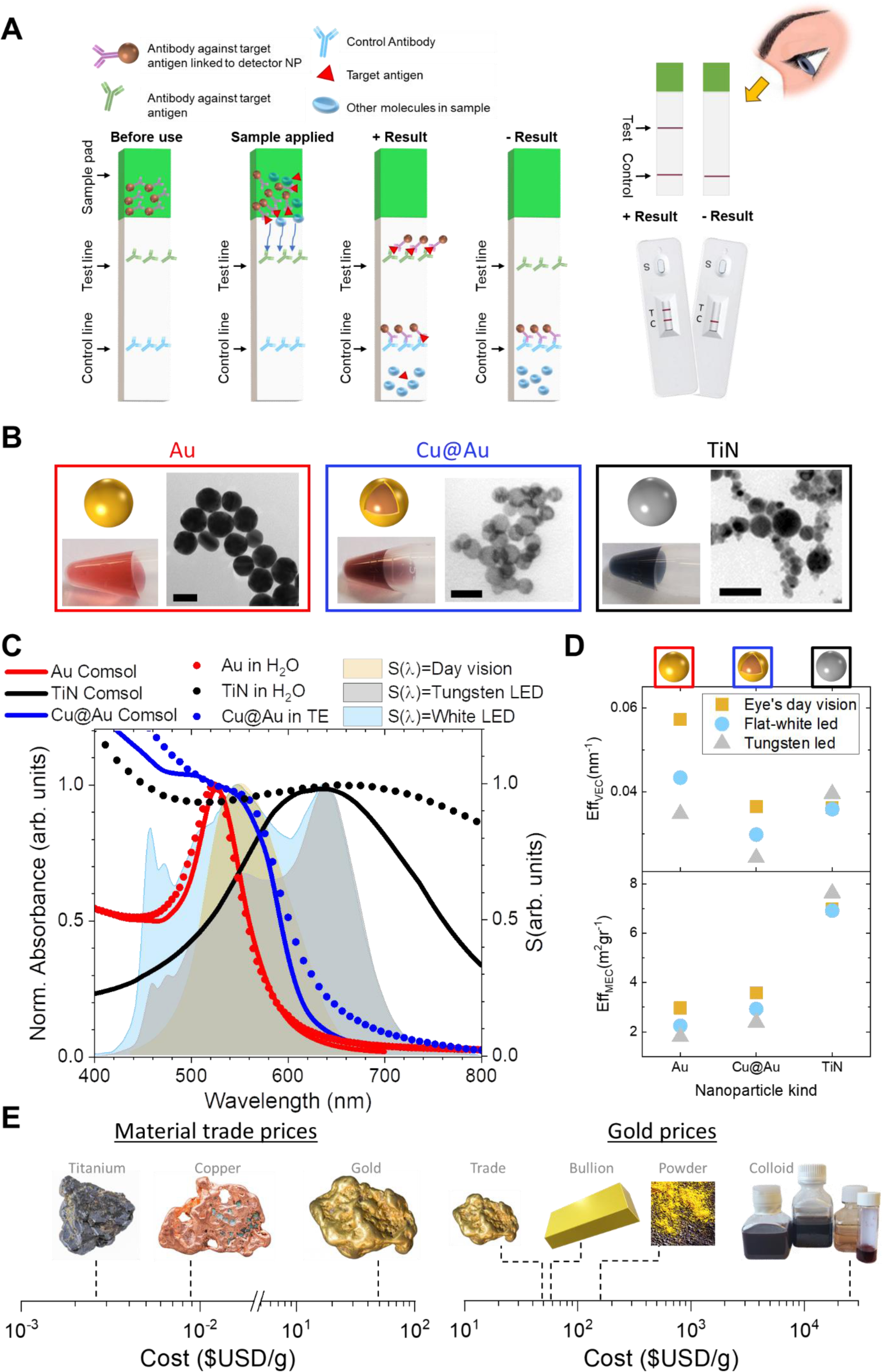
(a) Schematic representation of a lateral flow assay (LFA) showing the sample pad containing the marked antibody against the target antigen and the test and control lines. After the sample is applied to the sample pad, the sample plus the antibodies diffuse through the strip. Only if the target antigen is present, the marked antibody will bind to both the test and control lines (+ result). If the target antigen is not present, the target antibody will only bind to the control line (-result). The binding of the target antibody to the control line is necessary to validate a LFA test. (B) Characterization (scheme, colloid, and TEM) for the three kinds of NPs used throughout this work: Au in H_2_O [from ^35^], Cu@Au in Tris EDTA (TE) buffer, and TiN in H_2_O. Scale bars: 40 nm for Au, 20 nm for Cu@Au, and 100 nm for TiN. (C) Extinction cross sections from computational simulations for Au, Cu@Au, and TiN, all in a water environment (ε_NP_ =1.8) and each NP for diverse experimentally characterized sizes of 40 nm, 30 nm (28 nm TiN and 2 nm of TiO2 shell), and 12 nm (11.5 nm of Cu and 0.5 nm of Au shell), respectively. The shaded areas show three different assessment spectra: i) in light yellow, the day-vision sensitivity of the human eye (photopic vision), S_day−vision_ (λ); ii) in light gray, a typical tungsten LED; iii) in light blue, a typical white LED, S_white−LED_ (λ). (D) The VEC and the MEC efficiencies for each NP as shown, for the three spectral sources from panel C. (E) Left panel: Bulk material trade prices for titanium,^36^ copper^37^ and gold;^38^ Right panel: Sale prices for different gold products: trade, bullion, powder, and colloids [from Ref. ^13^].

Here we focus on the biofunctionalization and subsequent LFA implementation with spherical copper nanoparticles covered with a gold shell (Cu@Au) and spherical titanium nitride (TiN) NPs, as they have been proved to be stable, bio-friendly and capable of being manufactured at large scale.^17–20^ The choice of these NPs would be especially profitable when they are produced by cost-efficient and scalable methods such as laser ablation, which is much cheaper than colloidal chemistry under high production rates (> 500 mg/hour).^13^ TiN NPs show a plasmonic peak in the range of 650-800 nm and high photothermal conversion efficiency.^16,21^ TiN biomedical application has been limited by the difficulty of fabricating large-scale spherical-shaped in water. We recently demonstrated femtosecond laser ablative synthesis of large amounts of high quality TiN NPs,^18,20,22^ with already proven bioefficiency and low toxicity *in vitro* and *in vivo*,^18,19^ photothermal therapy on U87–MG cancer cell cultures under near-infrared laser excitation,^18^ and photoacoustic biological imaging.^22^ In general, laser-synthesized TiN NPs present a safe choice for future biomedical applications.^23,24^ Moreover, ultrastable Cu@Au NPs, have been recently synthetized via a seed-mediated galvanic replacement approach, which unlike CuAu alloys, the thin Au shell here provides enhanced stability. Furthermore, they show a superior photothermal efficiency in solar-induced water evaporation as compared to Au NPs.^17^ In general, TiN and Cu-Au NPs are promising to be useful in several fields such as catalysis, light harvesting, optoelectronics, and biotechnologies.^25–30^

We assess the biotesting LFA-performance of three different NPs (Fig. 1B) and show that inexpensive plasmonic TiN- and Cu@Au-based LFAs perform comparable to Au NPs-based LFAs, in some cases even better. Firstly, NPs were transferred from their synthetization organic solvents to biofriendly solvents, such as water and Tris-EDTA buffer. Then, the performance of antibody-functionalized TiN and Cu@Au NPs for detecting two kinds of biomolecules, Fluorescein isothiocyanate (FITC) and cardiac troponin (cTnT), was characterized on LFA strips. Our results show that the binding of the functionalized Cu@Au and TiN NPs allows an easy visualization of the target protein by naked eye, providing a signal efficiency and specificity similar to the observed when using the Au NPs. We further confirm these signals with machine learning (ML) techniques. Our approach is a starting point for the massive usage of these inexpensive NPs in bio-applications, in where the NPs’ plasmonic features in combination with their successful functionalization capabilities, may be useful in photothermal therapies, delivery, imaging and sensing technologies.

## Results

### COMSOL Multiphysics® simulations

COMSOL Multiphysics® numerical simulations were carried out to assess and quantify the optical performance of our three NPs. See full details in the Supporting Information.^31–34^ We simulate NPs illumination with linearly-polarized light (Eqs. S1-S3) and calculate the system’s optical extinction, which is the counterpart of the experimental absorbance measurements. The extinction was fitted to the absorbance, finding excellent agreement, which was later used to calculate the efficiency parameters for volume extinction coefficient (VEC) and mass extinction coefficient (MEC). The anticipated optical performance of these NPs in the presence of an excitation/sensing spectrum *S*(*λ*) can be estimated by the VEC and MEC magnitudes given by Eq. S4. Here we calculate four different *S*(*λ*) under which LFAs may be assessed: the human eye’s day vision, a tungsten LED, a white LED (Fig. 1C-D), and a cold cathode fluorescent lamp (Fig. S2). These quantities indeed show similar or even better magnitudes for TiN and Cu@Au NPs, anticipating that both NPs should perform as good as Au NPs.

### Cu@Au and TiN NPs preserved their characteristics after being transferred to an aqueous solvent

The Cu@Au NPs synthesized via galvanic replacement, were dispersed in hexane, containing an oleylamine coating layer. Wang *et al* described that these ∼12 nm particles showed high stability even in harsh environment since the gold shell completely protects the copper core.^17^ The TiN NPs were synthesized via laser ablation, allowing the production of large amount of spherical size-specific NPs, which do not aggregate due to electrostatic stabilization. These NPs of ∼30-40 nm were grown in acetone, showing great stability and capability of by polyethylene glycol-coating.^18,20,22^ However, to be used in biological devices, it is more convenient to preserve these NPs in aqueous solvent but maintaining their original characteristics. First, we successfully transferred the Cu@Au and TiN NPs to aqueous media by ligand-exchange and evaporation of acetone, respectively (Figure S4). The characterization of the Cu@Au NPs indicates that they maintain an average size of 12 nm (Weibull fit, max. 11.3 nm) when suspended in the aqueous medium (Fig. S4A). Their observed plasmon was around 555 nm, which corresponds to their original plasmon observed when suspended in hexane (Fig. S4B). The stability analysis in the aqueous medium after measuring the plasmon at day zero and >40 days, indicates that the stability of the Cu@Au NPs is preserved (Fig. S4C). The characterization of the TiN NPs indicates these particles have an average size of 35 nm (Weibull fit, max. 34.5 nm) when suspended in water (Fig. S4D). The observed plasmon was around 650 nm, in good agreement with the observed plasmon before their transfer to water (Fig. S4E). TiN NPs were also stable through time since their plasmon was preserved after >40 days (Fig. S4F). Together, these results show that after transferring Cu@Au and TiN NPs to aqueous media, the NPs size and optical properties were preserved.

### Functionalized Cu@Au and TiN NPs showed similar efficiency to Au NPs in LFA

To determine if these NPs could be used as biosensors, we tested their capability to detect an antigen of interest. For the analysis, Cu@Au and TiN NPs were functionalized with an antibody anti-FITC. As a control, commercial 40 nm Au NPs (NanoComposix®) were used and tested simultaneously. As shown in Figure 2A, after selection of the appropriate buffer and functionalization, the control Au, the Cu@Au, and TiN NPs were maintained in homogeneous suspension. To corroborate functionalization, absorbance was measured before and after functionalization. Figure 2B shows a plasmon red-shift for all the NPs, suggesting a few nm shell of new dielectric media (in our case, biomolecules), evidencing a successful modification step. Moreover, the efficiency of the functionalization was similar for the three NPs, as demonstrated by their absorbances (Fig. 2B). For testing the ability of the functionalized NPs to detect the antigen of interest, LFA were run. Figure 2C shows the functionalized TiN and Cu@Au NPs identified the target protein (FITC-T10-Biotin) present at a concentration between 100 and 1 nM. We quantified the intensity of each LFA band for the three NPs, determining that the intensity of the signal given for the TiN NPs was similar to the intensity registered when using the Au NPs (Fig. 2D). The intensity of the LFA bands when using the Cu@Au NPs (∼12 nm) was ∼37% lower compared to the Au NPs (Fig. 2D). However, considering the size of the tested particles, our results indicate that both NPs, Cu@Au, and TiN, can be successfully functionalized and effectively detect a specific target protein with similar or even better efficiency to the well-standardized Au NPs. The NPs’ plasmonic characteristics were preserved even when attached to the test and control line of the LFA strip, as demonstrated by their ability to absorb at specific wavelength (see Figure S5).

**Figure 2.**
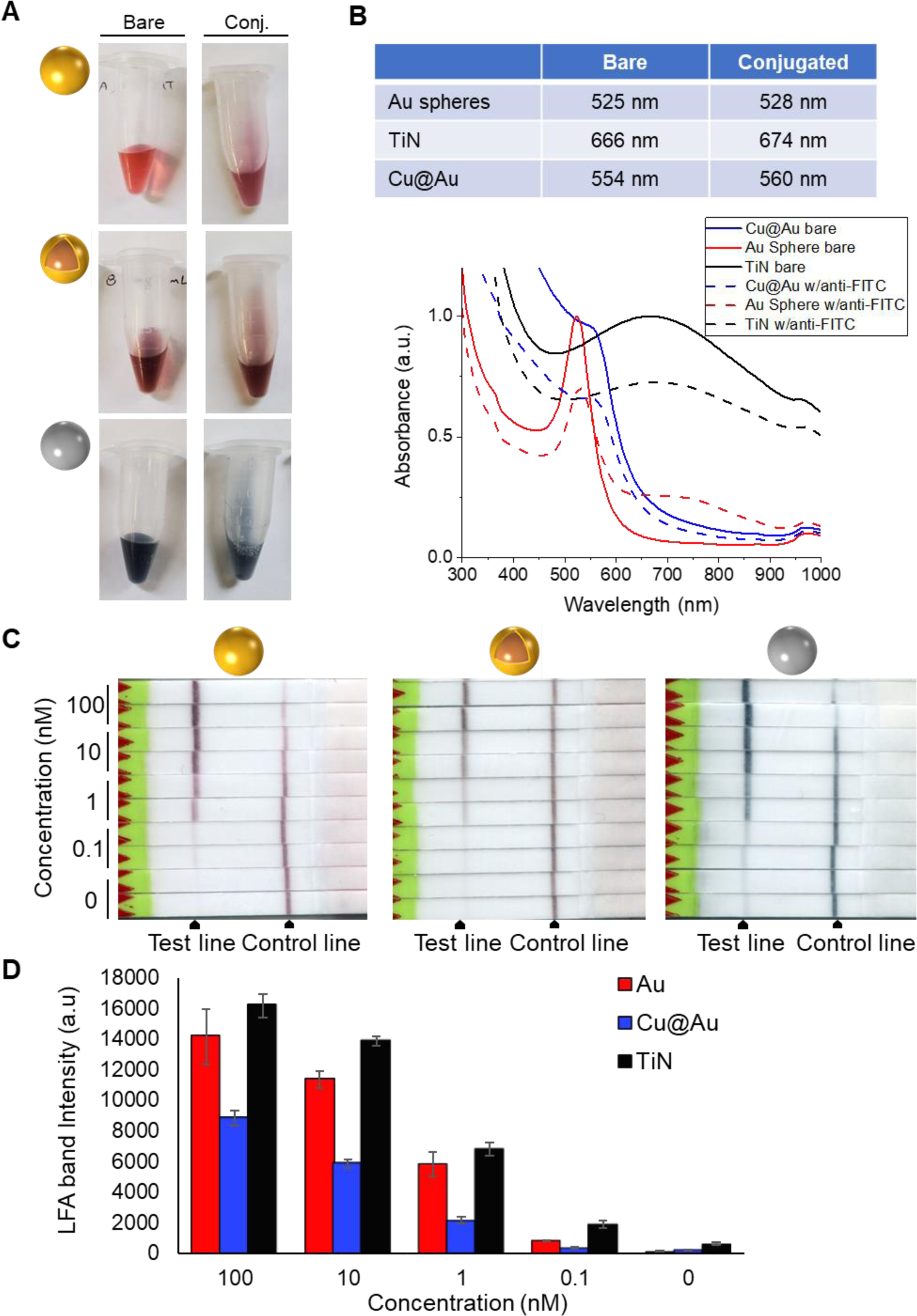
Functionalization of NPs with anti-FITC antibody. (A) Visual characterization of each type of NPs before and after antibody conjugation to demonstrate they do not aggregate nor precipitate. (B) Table and plot comparing the plasmons for each type of NPs before (solid lines) and after (dashed lines) conjugation. (C) LFA in duplicate showing the identification of FITC at different concentrations (100, 10, 1, 0.1, and 0 nM) using the conjugated NPs. (D) Plot showing the intensity measured via ImageJ of each test band of the LFA.

### Functionalized Cu@Au and TiN NPs can detect proteins with biological significance

Since our first functionalization analysis involved the recognition of the system FITC-T10-Biotin, we tested if the particles can detect proteins with biological significance. Here, NPs were functionalized for detecting cardiac troponin (cTnT), a protein that increases in the blood of patients suffering a heart attack. The Au (positive control), Cu@Au and TiN NPs were modified with an antibody anti-cTnT. Figure 3A shows a change in the migration pattern in a 0.25% agarose gel, confirming the functionalization of the three NPs. Additionally, the visual analysis (Fig. 3A) and TEM imaging (see Figure S6A) of the NPs solutions confirm that the NPs do not aggregate and remain mono-dispersed. In addition, a red-shift in the Cu@Au and TiN NPs absorbance pattern (Figure S7) further confirms their successful functionalization (note that in Fig. S7 the post-functionalization absorbances have been normalized to facilitate comparison). Next, different concentrations of recombinant human cTnT were blotted in a nitrocellulose membrane as schematically shown in Figure 3B. After blotting, the presence of the proteins in the nitrocellulose membrane was confirmed by Ponceau S staining (Fig. 3C, left). The membrane was then divided into three equal pieces, incubating each of them with one of the NPs. Figure 3C (right), confirms that our three conjugated NPs were able to bind to the target protein (cTnT), displaying a signal that correlates with the amount of protein blotted in the membranes. The signal was observed quickly, after 10 minutes of incubation with each NP. These results confirm that the TiN and Cu@Au NPs can be functionalized with different antibodies to identify proteins with biological significance. Finally, to test the specific recognition of the target protein, LFA was developed as displayed in Figure 3B, using bovine calf serum with (+) or without (-) cTnT (0.03 ug - 0.04 ug). Each LFA confirms the specificity of the test since the test bands were only observed in the presence of cTnT (Fig. 3D). The binding of the functionalized Cu@Au and TiN NPs allow an easy visualization by the naked-eye, with a signal efficiency and specificity similar to the observed when using the Au NPs (Fig. 3C-D).

**Figure 3.**
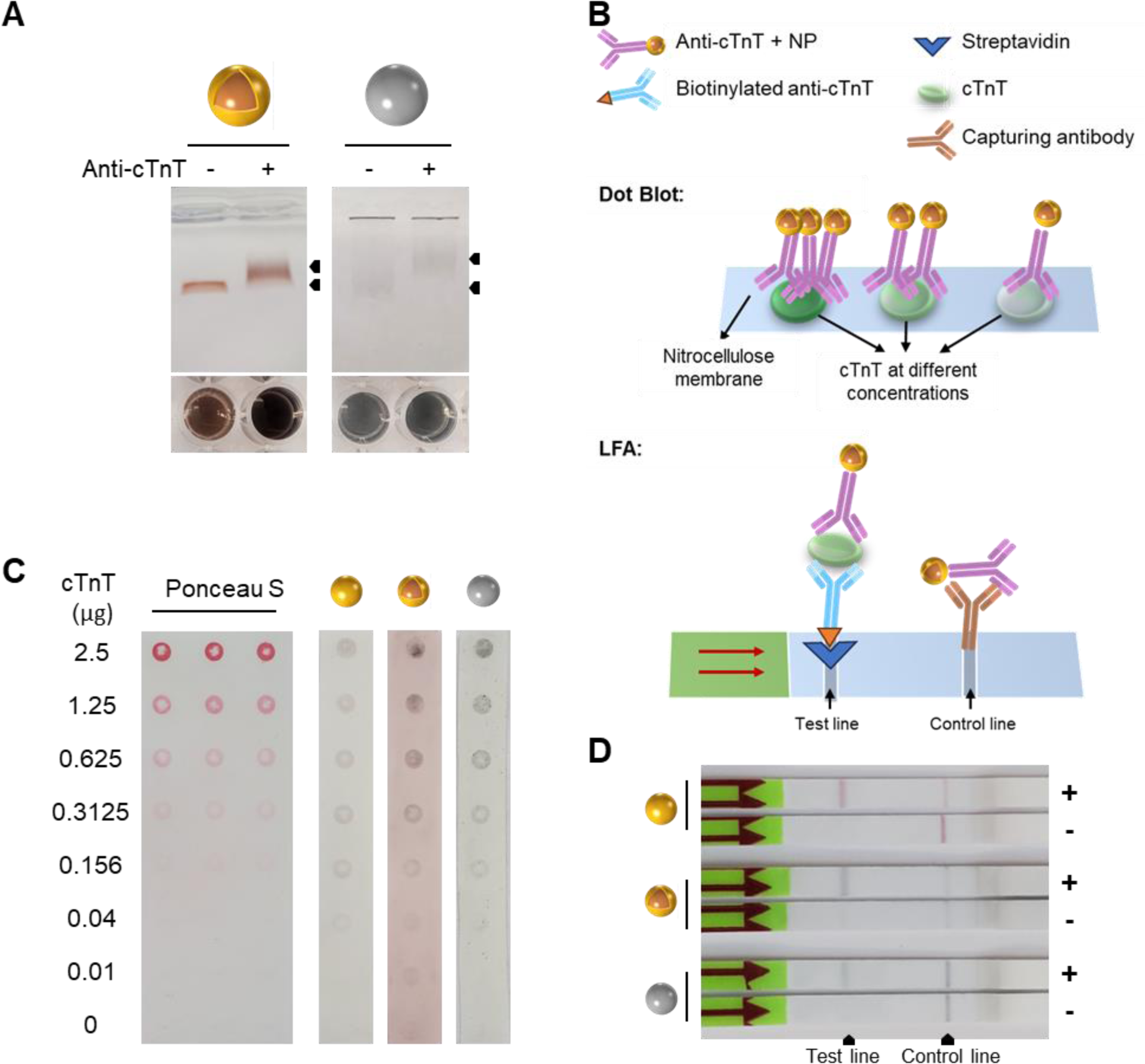
Using Cu@Au and TiN functionalized NPs for naked-eye detection of cardiac troponin. (A) 0.25 % agarose gel for testing the functionalization of the Cu@Au and TiN NPs. At the bottom of the gel, each solution pre- and post-functionalization is shown. (B) Representative schemes of the dot blot (top) and LFA (bottom) used for testing, respectively, the ability of the Cu@Au and TiN NPs to detect the target protein and their specificity. (C) Dot blot analysis showing the detection of cardiac troponin (cTnT) at different concentrations (µg). The Ponceau S staining corroborates the presence of cTnT at different concentrations in the nitrocellulose membrane. The membrane was cut into 3 sections, which are shown after being incubated for a total of 30 minutes with the conjugated Au, Cu@Au, and TiN NPs. (D) LFA against cTnT diluted in bovine calf serum, showing the specific detection of cTnT when using each conjugated NP.

### Common machine learning platform and experiments to evaluate Cu@Au and TiN-based LFAs in the detection of FITC

We aim to establish a common and fair ML platform for comparing the discriminative capabilities of the proposed TiN and Cu@Au-based LFAs against Au-based LFAs. Toward this objective, we developed two methods: 1) feature extraction based on the Red-Green-Blue (RGB) color channels of the test and control lines on the strips; 2) contrastive learning, an advanced ML model, to enhance feature extraction. The K-Nearest Neighbor (K-NN) classifier is utilized in both methods to perform strip classification.

In our experimental dataset, pairs of strips composed of TiN, Cu@Au, and Au were tested with six different FITC concentrations. We manually segmented the test and control lines on each strip, focusing on their central areas (Fig. 4A). The Red/Green/Blue (R/G/B) channels of all lines were fitted with Gaussian distributions, from which we extracted the mean, along with the lower and upper bounds of the first standard deviation. Each line is represented as a nine-element vector, and each strip’s representation is an 18-element vector, formed by concatenating the vectors of the test and control lines.

**Figure 4.**
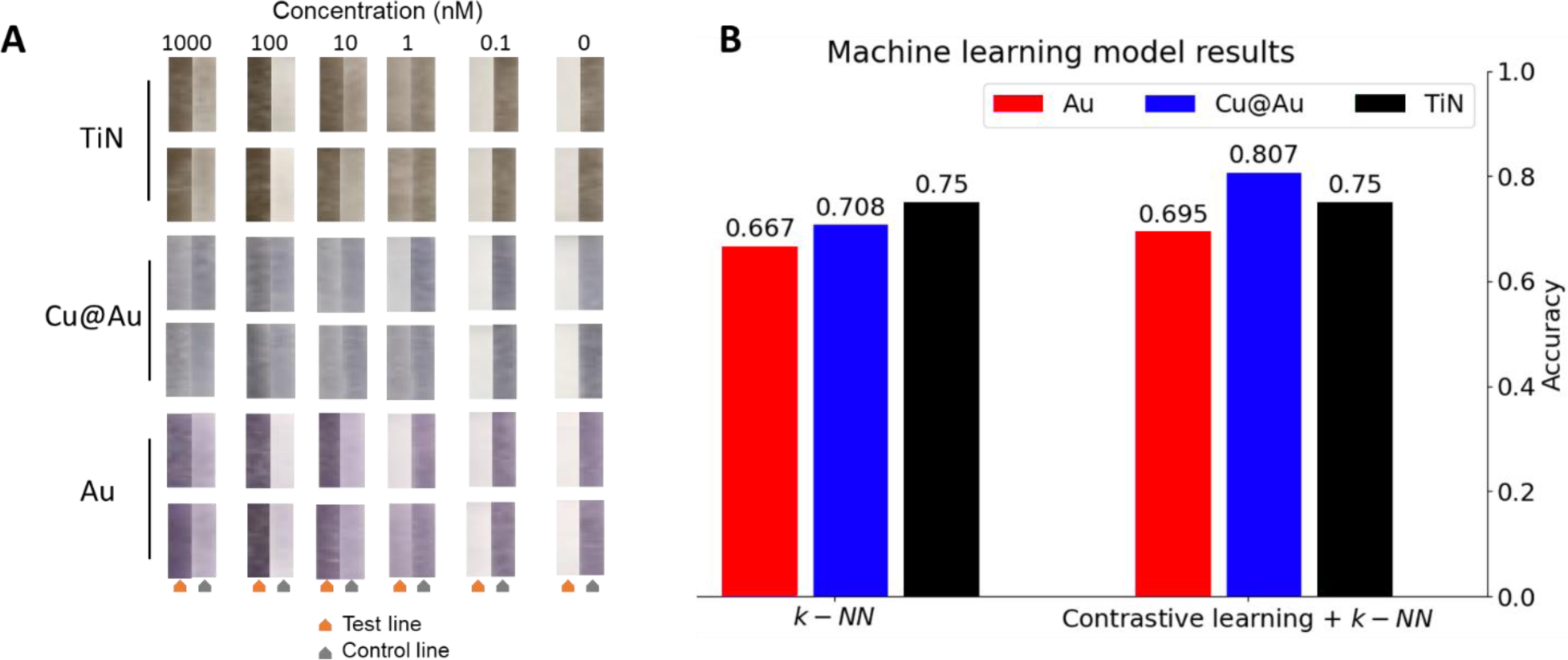
Data, models, and results of our machine learning experiments. (A) Cropped test lines and control lines of all strips that tested with different FITC concentrations. (B) Results of the machine learning experiments. It is evident that the accuracy of TiN and Cu@Au-based LFAs surpasses that of Au-based LFAs.

Figure 4B shows the accuracy results for the different NPs with the k-NN model, indicating that TiN and Cu@Au achieve higher accuracy than Au. This can be considered evidence that TiN and Cu@Au LFAs are more robust and responsive in detecting FITC concentrations.

Moreover, we developed an advanced feature extraction approach known as contrastive learning to further explore the NPs discriminative capability. Following training, the contrastive model is employed to generate representation vectors for the input data. We then apply the k-NN classification method on the vectors, using the same experimental setup as in our previous experiment (Fig. 4B). Through the application of contrastive learning,^39^ both Cu@Au and Au demonstrated improved performance, while TiN’s performance remained unchanged. However, the accuracy for TiN and Cu@Au still surpassed that of Au, indicating that the features of TiN and Cu@Au-based LFAs are more distinguishable.

## Discussion

The COVID-19 pandemic showed the utility of the easy-to-develop and affordable LFA tests. These tests are made mostly of gold, a noble metal with great chemical stability and strong plasmonic absorption. However, price and global availability are some issues that can affect its massive use in close future ^14^. In this study, we proved that TiN and Cu@Au NPs-based LFAs perform comparable to Au NPs-based LFAs. We firstly show that these novel inexpensive NPs can be successfully transferred to bio-friendly (aqueous) medium for later biological uses (Fig. S4B-E), showing stability as a function of time (Fig. S4C-F), so they could be eventually used in long-times delivery operations. We then show that both NPs, can be successfully functionalized with diverse antibodies, demonstrated by the plasmon red-shifts and agarose gel analysis (Figs. 2 and 3). Furthermore, the antibodies conjugated to both NPs preserve their function, confirmed by the identification of different concentrations of FITC (Fig. 2) and cTnT (Fig. 3).

Antibody orientation onto NPs is an important factor broadly studied and discussed^40–43^ since their proper orientation on the NPs’ surface, preserves their ability to specifically detect the target molecule. Au NPs have demonstrated this ability, which is also shown in this study for the TiN and Cu@Au NPs. The conjugation of TiN NPs, as well as for Au control NPs, involved the non-covalent immobilization of the target antibody onto their surfaces. However, this did not work with the Cu@Au NPs due to the presence of the carboxyl groups from the 3-MPA. In this study we included EDC, which reacts with the carboxyl group of the 3-MPA, followed by the incorporation of a hetero-bifunctional crosslinker containing an amine-reactive N-hydroxysuccinimide (NHS) ester and a photoactivatable nitrophenyl azide, to ensure the covalent interaction between the Cu@Au NP and the target antibody. Additionally, TiN NPs conjugation with DNA via instant dehydration in butanol^44^ was tested (Figure S6B), demonstrating the versatility and stability of their conjugation. By the successful conjugation and identification of two different markers, our results prove that the TiN and Cu@Au NPs allow a working flexibility comparable to Au NPs. This working flexibility is an important characteristic since indicates that these inexpensive NPs can be broadly used in biotesting technologies. Additionally, when the conjugated NPs were exposed to a sample containing a mix of proteins (cTnT and bovine calf serum), the specific detection of cTnT (Figure 3) confirms that the TiN and Cu@Au NPs have similar properties compared to the Au NPs.

We also demonstrated that the signal provided by these novel NPs at naked-eye is comparable to the Au NP’s signal, as it was qualitatively predicted by our simulations (Fig. 1C). The NPs’ optical efficiencies of typical assessment sources were calculated (Eq. S4), estimating how “seeable” the NPs-loaded LFA lines will be. We find that Eff_*VEC*_ correlates well with our measured experimental intensities (Figures 1D versus 2D), when using the tungsten LED. The experimental measured lower signal of the Cu@Au NPs-loaded LFAs could be explained by the smaller size of these NPs, since the surface area of a single Cu@Au NP was about 90% less than that of a single Au and TiN NP. We expect that if we use same mass of NPs, Cu@Au NPs can outperform Au NPs. Also, the experimental higher signal of the TiN NPs-loaded LFAs could be explained by the polydispersity of larger size of these NPs (Fig. S4D). Furthermore, dot blot and LFA (Figs. 2-3) demonstrate that the observed signal correlates with the amount of target protein, suggesting that after a suitable numerical preparation and subsequent experimental standardization, this technique can be also applied for a quantitative analysis.

We should also comment on other inexpensive materials for LFAs, such as latex, polystyrene and carbon, among others. However, several of these devices require sophisticated equipment for reading out the results, hence delaying a possible diagnosis.^45–48^ Additionally, these materials do not show the benefits of plasmonic systems, such as propagating/localized surface plasmon resonance (SPR), surface-enhanced Raman scattering (SERS), fluorescence and infrared absorption spectroscopy, among others,^49^ broadening the possible applications in where our NPs could be used.

As for our ML approach, the simple k-NN algorithm obtains an accuracy higher than 70% for TiN and Cu@Au-based LFAs. Through contrastive learning, the accuracy surpasses 75%. The visual features exhibited by TiN and Cu@Au-based LFAs prove to be robust, responsive, and consistent in detecting various FITC concentrations, indicating their potential as viable alternatives to Au-based LFAs with broader applicability in bio-related domains. Also, the anticipated Eff_*MEC*_ correlates well with our k-NN algorithm (Fig. 1D versus 4D). Moreover, the accuracy obtained in our ML experiments, suggests the potential for automated quantitative analysis of biomolecules using TiN and Cu@Au-based LFAs.

In conclusion, we showed here a comprehensive numerical-experimental study demonstrating that the performance of our inexpensive NPs for detecting FITC and cTnT, was similar to the performance of the current gold standard Au NPs. For the ML approach, the TiN NPs show an advantage in recognition accuracy that speaks to the potential of new materials in biosensing.

## Acknowledgements

V.A.B.L. and S.W. acknowledge the partial support by NIH ES030425 (to S. W.) and the Edison Biotechnology Institute, Ohio University (V.A.B.L. and S.W.). O.A.O. and A.O.G. acknowledge the generous financial support from the Baker Fund, QBI, and NQPI at Ohio University. E.S. and K.L.A.C. acknowledge the Department of Chemistry and Biochemistry and the College of Arts and Sciences. D.M. is grateful to the support of Canada Research Chairs Program (CRC 950-232964), Natural Sciences and Engineering Research Council of Canada (RGPIN-2020-05921) and the Fonds de recherche du Quebec-Nature et technologies.

## Author contribution

**Veronica A. Bahamondes Lorca**: Bioconjugation of NPs, LFA testing, NP’s ligand transfer, conceptualization, and manuscript writing, **Oscar Ávalos-Ovando**: Bioconjugation of NPs, LFA testing, simulations, conceptualization, and manuscript writing, **Christoph Sikeler**: Bioconjugation of NPs, LFA testing, conceptualization, imaging, manuscript writing, **Heini Ijäs**: Bioconjugation of NPs, and LFA testing, **Eva Yazmin Santiago**: simulations, **Eli Skelton**: UVvis data acquisition, **Yong Wang**: NP’s synthesis, and ligand transfer, **Ruiqi Yang**: NP’s synthesis, and ligand transfer, **Katherine Leslee A. Cimatu**: Work supervision, **Olga Baturina**: Optical experiments with plasmonic strips, **Zhewei Wang**: Machine learning, and manuscript writing, **Jundong Liu**: Machine learning, and manuscript writing, **Joseph M. Slocik**: Conjugation protocol, and data discussion, **Shiyong Wu**: Work supervision, **Dongling Ma**: NPs synthesis, work supervision, data discussion, and manuscript writing, **Andrei I. Pastukhov**: NPs synthesis, **Andrei V. Kabashin**: NPs synthesis, work supervision, data discussion, and manuscript writing, **Martin E. Kordesch**: Imaging, work supervision, and data discussion, and **Alexander O. Govorov**: Conceptualization, manuscript writing, data discussion, and work supervision.

## Supporting Information for

### I. Materials and Methods

#### Synthesis of Cu@Au NPs

Typically, 33.2 mg of copper(II) acetylacetonate (Cu(acac)2, ≥99.9%, Sigma-Aldrich) was dissolved in 10 mL of Oleylamine (OLA, 70%, Sigma-Aldrich) under nitrogen atmosphere. Then, the solution was kept at 230 °C for 3 h to produce Cu NPs. Subsequently, the solution was cooled down to 140 °C and then 1 mL of Trioctylphosphine (TOP, 90%, Fisher Scientific) solution of gold(III) chloride trihydrate (HAuCl4·3H2O, ≥49.0%, Sigma-Aldrich) (20 mg) was injected. After 1 h reaction, the solution was quickly cooled down to room temperature. Finally, the Cu@Au NPs were purified and dispersed in hexane for further use.

#### Cu@Au NPs ligand exchange

1 mL of Cu@Au dispersion in hexane was added to 2.5 mL of methanol. The solution was alkalinized by adding 140 μL of NH4OH (Sigma). Then, dropwise, 90 μL of 3-Mercaptopropionic acid (3-MPA, Sigma) was added. The solution was heated in a hot plate at 75°C for 1h with manual sporadic agitation to yield the 3-MPA capped Cu@Au NPs. The NPs were then precipitated by adding 10 mL of distilled water, 2.5 mL of isopropanol, and 1% Triton x100. The Cu@Au NPs were collected by centrifugation at 7000 rpm for 10 min, rinsed with water, and resuspended in 1X Tris-EDTA (TE), pH 8.0 buffer.

#### TiN NPs synthesis

TiN NPs were synthesized by the technique of femtosecond laser ablation in liquid ambient. A TiN target (MaTeck, Germany, 99+%) was fixed in a vertical position inside a glass cuvette (Hellma, Germany, optical glass, 88 mL, 2.5 mm wall thickness) filled with 80 mL of acetone (Acros Organics, 99.5+%). The thickness of the liquid layer between the target and a cuvette wall was 3 mm. To initiate material ablation, a femtosecond laser beam (s-Pulse HP, Amplitude Systems, France, Yb:KGW, 490 fs, 10 kHz) was focused through the cuvette wall on the surface of the target using a 75 mm convex lens. The energy was attenuated down to 150 µJ per pulse using a half-wave plate and Brewster polarizer. To prevent ablation from the same area, the target was constantly moved at a speed of 2.5 mm s-1 by a translation stage (scanned area was 5 x 5 mm2).

#### TiN NPs water dispersion

Distilled water was added to a TiN dispersion in acetone in a ratio of 1:1. The mixture was heated on a hot plate at ∼150°C under constant agitation until all the acetone was evaporated from the sample.

#### Electron microscopy

NPs drop-cast from solution onto Lacey Carbon TEM grids to facilitate various types of NP characterization. A JEOL 1010 Transmission Electron Microscope (TEM) operated at 80 kV and connected to a MSC TK1024M camera (Gatan, Munich, Germany), was used to obtain the NPs morphological details.

#### Functionalization of NPs with FITC

NPs were centrifuged and dispersed in their corresponding buffer (Au: TE pH 8.0, Cu@Au: Borate pH 7.0, and TiN: 1X TE pH 8.48). 2 μg of anti-FITC (200-032-037, Jackson ImmunoResearch Laboratories, Inc.) per 40 μL of NPs solution was added and incubated in a rotor at 4°C for 30 min. 10% BSA was added to the NPs to have a BSA final concentration equal to 1 %. The samples were incubated for an extra hour at 4°C with constant rotation. The NPs were then rinsed twice and centrifuged at 13,000 rpm for 10 min each. Finally, the NPs were resuspended in their corresponding buffers.

#### Functionalization of NPs with anti-cTnT

Au and TiN NPs were centrifuged and then dispersed via sonication in their corresponding buffer (Au: 1X TE pH 8.0 and TiN: 1X TE pH 8.48). 10 μL/mL of Anti-Cardiac Troponin T antibody [1F11] (cTnT, ab10214, Abcam) was added to each sample and incubated in a rotor at room temperature for 30 min. 10% BSA was added to the NPs to obtain a BSA final concentration equal to 1 %. The samples were incubated for an extra hour at room temperature with constant rotation. The NPs were then rinsed twice and centrifuged at 4 °C and 12,000 rpm for 10 min each. Finally, the NPs were resuspended in their corresponding buffers. For the Cu@Au NPs, after centrifugation, NPs were resuspended in 100 mM MES buffer pH 6.0 containing excess of freshly prepared 1-Ethyl-3-[3-dimethylaminopropyl]carbodiimide hydrochloride (EDC, 1 mg/mL, # 22980, Thermo Scientific™). Excess of sulpho-sulfosuccinimidyl 6-(4’-azido-2’-nitrophenylamino)hexanoate (sulpho-SANPAH, 1 mg/mL, # 22589, Thermo Scientific™) was added immediately to the NPs solution and samples were incubated in dark, at room temperature and under constant rotation for 20 min. 2-Mercaptoethanol (10 μL/mL) was added to the NPs to quench EDC, samples were incubated under the same condition for an additional 20 min. Same volume of 100 mM phosphate buffer pH 7.5 was added to increase the pH of the Cu@Au NPs, following for 10 μL/mL of anti-cTnT antibody (ab10214, Abcam). Samples were transferred to a flat container and irradiated with UV from a 340-UVA lamp (Q-lab, Co.) for 45 minutes at 1.15 mW/cm2. NPs were centrifuged at 12,000 rpm and 4 °C for 10 minutes and rinsed in 1X TE buffer pH 8.0.

#### Agarose gel separation

0.25 % agarose gel was prepared in 0.5 X Tris-borate-EDTA (TBE) buffer (40 mM Tris-base, 40 mM Boric acid, and 1 mM EDTA). The samples were loaded with sucrose 50 % and run at 100 Volts for 5 to 10 min.

#### Lateral flow assay (LFA)

For FITC-LFA, 20 μL of anti-FITC functionalized NPs were mixed with 0.2 μL of the oligo FITC-T10-Biotin (Metabion International AG, Germany) diluted at the specified concentrations. Commercial LFA strips (MGHD1, Milenia GenLineHybridetect, Germany) were used, which were produced at request without the gold NPs pad. The samples were run following the manufacturer’s indications. The strips were scanned and the intensity of each band in the test strips was quantified using Image J. For cTnT LFA, commercial LFA strips (MGHD1, Milenia GenLineHybridetect, Germany) were loaded with 4 μL (Au/TiN) or 8 μL (Cu@Au) of conjugated NPs (OD ∼1.0). 10 μL (for Au/TiN LFA) or 13 μL (for Cu@Au LFA) of sample mix (50 μL of running buffer, 25 μL 1:1 bovine calf serum/PBS containing 0.156 μg of recombinant human cTnT (ab86685, abcam), and 1 μL of biotinylated anti-cTnT antibody [bs-10648R-Biotin, Bioss]) was loaded in the LFA strips. The sample mix for the negative controls only contains running buffer, biotinylated anti-cTnT antibody and bovine calf serum in PBS (1:1).

#### Dot blot

Recombinant human cTnT (ab86685, abcam) was diluted in 1X PBS buffer. 50 μL of the protein at different concentration were blotted onto a nitrocellulose membrane (0.2 μm, #88024, Thermo Scientific™). After aspiration was completed, the membranes were stained with Ponceau S to check the presence of the blotted protein on them. The membranes were rinsed with TBS-T buffer until completely removed the stain and then blocked in 5 % milk for 30 minutes. The membranes were rinsed once with each of the corresponding NPs-buffers followed by the incubation for a maximum of 30 min at 4°C with the conjugated NPs.

### II. Computational simulations

#### II.1 Comsol simulations

Our classical electromagnetic simulations on plasmonic nanoparticles are performed via the Finite Elements Method with the COMSOL Multiphysics simulations software. We illuminate the nanoparticle with linearly polarized light and calculate the optical response of the system. The incident electromagnetic field is defined as: 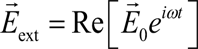. By solving the Maxwell’s equations within a classical framework, the calculations provide far-field quantities such the absorption (*σ*_*abs*_), scattering (*σ*_*scat*_), and extinction (*σ*_*ext*_) optical cross-sections (related by *σ*_*ext*_ = *σ*_*abs*_ + *σ*_*scat*_). The scattering cross-section is calculated by integrating the scattered intensity over a fictitious sphere around the NP, whereas the formalism for the absorption cross-section is based on the following equations:

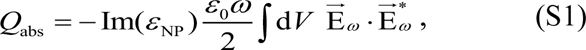

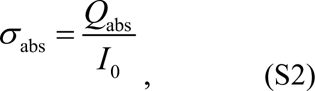

Where *Q*_abs_ is the absorbed power by the system, *ε*_NP_ is the dielectric constant of the metal nanoparticle, *ω* is the angular frequency of the incident light, 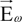 is the complex electric field inside the metal, and *I*_0_ is the photon flux magnitude (intensity for simplicity), given by

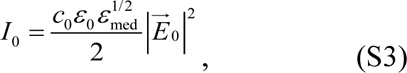

where *ε*_*med*_ is the dielectric constant of the medium, *c*_0_ is the speed of light in vacuum, *ε*_0_ is the vacuum permittivity, and 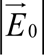 is the electric field magnitude of the incident electromagnetic wave. All NPs are simulated with sizes estimated from experimental characterization, as described in Fig. 1. In particular, for TiN NPs we find that a thin shell of TiO_2_ is needed in the accurate simulation of the extinction, as previously described elsewhere ^1^. For gold and copper we use the permittivity *ε*_NP_ from Johnson and Christy ^2^, for TiN from Guler ^3^, and for TiO_2_ from Siefke ^4^. The sum of *σ*_*scat*_ and *σ*_*abs*_ allows to obtain *σ*_*ext*_, related to the experimental measurement of absorbance.

#### II.2 Model for TiN NPs: TiN-TiO_2_ core-shell

Here we validate our particular choice of parameters for the core-shell model of TiN-TiO_2_ NPs.

For the spherical Au NPs and Cu@Au NPs the numerical simulations of extinction immediately fit the experimental absorption, as shown in Fig. 1C in the main text. This is done by using the experimentally measured dielectric constants available in the literature ^2^; using the typical TEM characterized sizes 40 nm for Au NPs and 12 nm (11.5 nm of Cu and 0.5 nm of Au shell) for Cu@Au NPs; and following the formalism shown in the Materials and Methods section S.I.

For the TiN NPs on the other hand, we need to account for oxidation and the inherent polydispersity inherited from the growth process ^1^. This is found to be only achievable when a core-shell of TiN-TiO_2_ is considered, accounting for slight oxidation of the outer NP shell. The TiN dielectric function is taken from Guler ^3^, and for TiO_2_ from Siefke ^4^, and several combinations of values for the TiN-core and TiO_2_-shell are tested. See Figure S1A for schematics. By simultaneously comparing with our experimental absorbance (red symbols), one can see that the total diameter of the NP does not play much role, but it just red-shifts the plasmon a few nanometers when simulating D=10 nm to D=40 nm (Figure S1). The TiO_2_ thickness on the other hand, does play a role by making the plasmon slightly wider and red-shifting it significantly, as shown in Figure S1C. Finally, these are the spectra (specifically, the one of 30 nm, 28 nm TiN and 2 nm of TiO_2_ shell) we used in Figure 1C and for calculating the efficiency parameters of Figs. 1D and S2.

**Figure S1.**
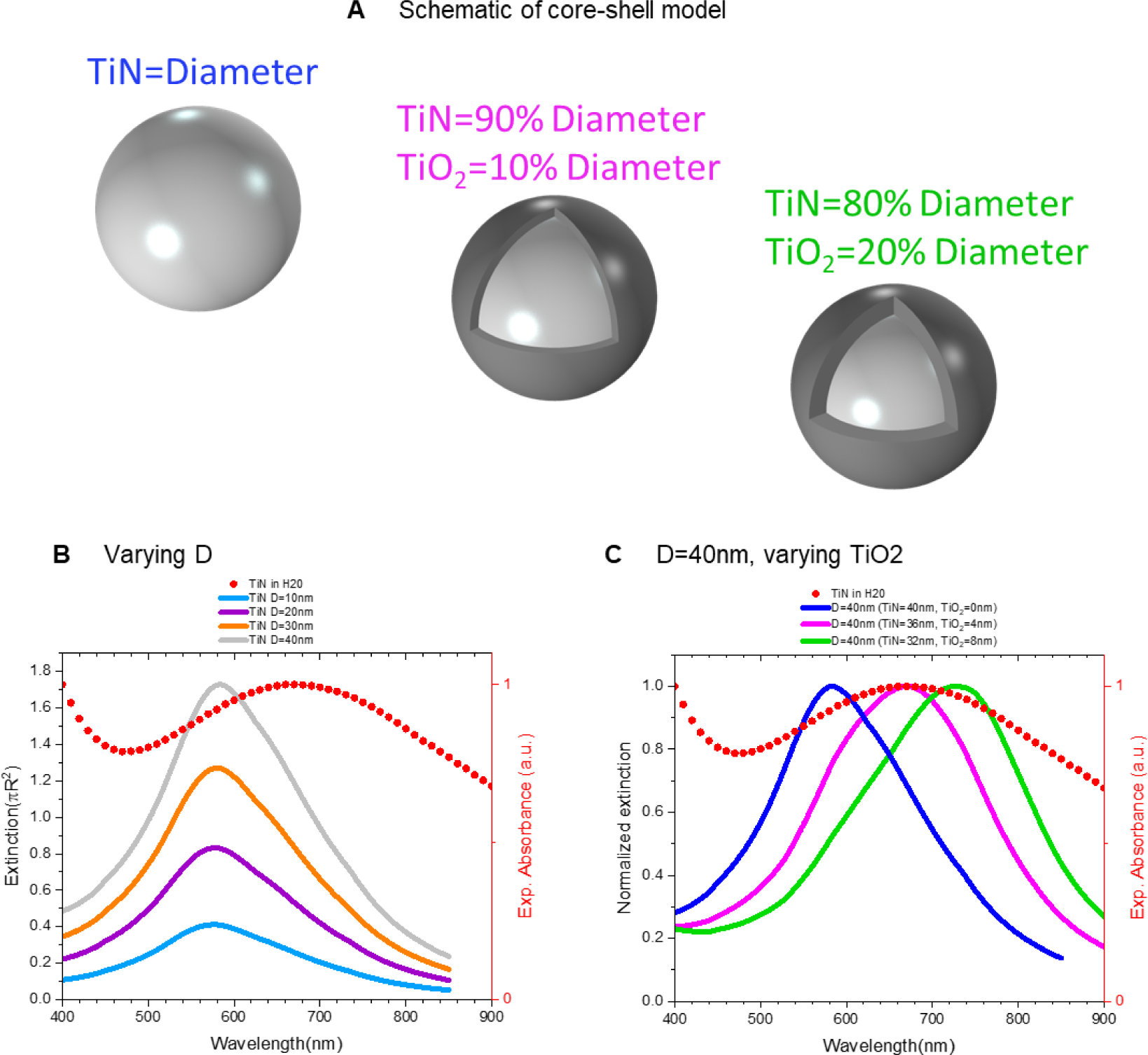
Models of TiN-TiO_2_ core-shell. (A) Schematic representation of different aspect ratios between TiN-TiO_2_ as shown. In all plots, lines are Comsol numerical simulations (left axis) and red symbols are our NPs experimental absorbance (right axis). (B) Different diameters of TiN, no TiO_2_. (C) Same NPs diameters (D=40nm) with three TiN-TiO_2_ aspect ratios.

#### II.3 Efficiency parameters

We also calculate the visual efficiency of each NP under the excitation of several common light sources and/or sensorial capacities, each of which will have a given signature spectra *S*(*λ*). The absolute efficiency is given by

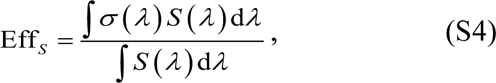

which allows us to quantify each NP performance on a visual assessment under such spectral conditions. Briefly, the calculation starts by fitting numerical extinctions with our experimental absorbances, from where then the first were used in Eq. S4, which is a convolution between *S*(*λ*) (shaded-regions color in Fig. 1C) and each NPs’ individual absorbance (plasmons in Fig. 1C), yielding the efficiencies shown in Fig. S2 and Fig. 1D. In order to perform a fair comparison, we then also normalize each NP’s extinction by the amount of material present, meaning by the NP’s volume and the NP’s mass in order to obtain the volume extinction coefficient (*σ*_*VEC*_ = *σ*_*ext*_/*volume_NP_*) and the mass extinction coefficient (*σ*_*MEC*_ = *σ*_*ext*_/*mass_NP_*), for then calculate the relative efficiencies Eff_*VEC*_ and Eff_*MEC*_. In particular, we use the volumes for the sizes described in Fig. 1 and the densities from Table S1.

**Table S1.**
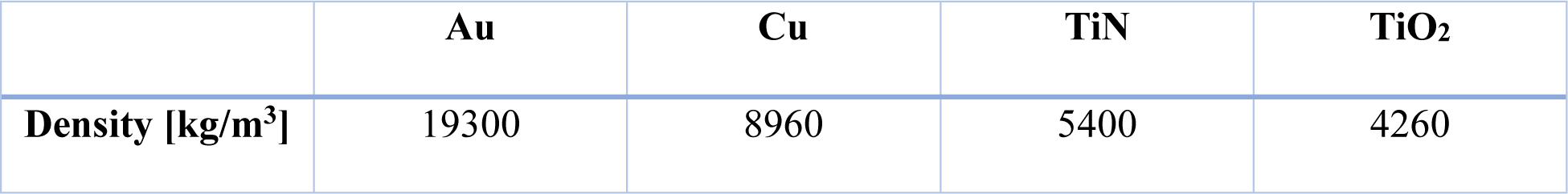
Densities for the materials used in our work.

**Figure S2.**
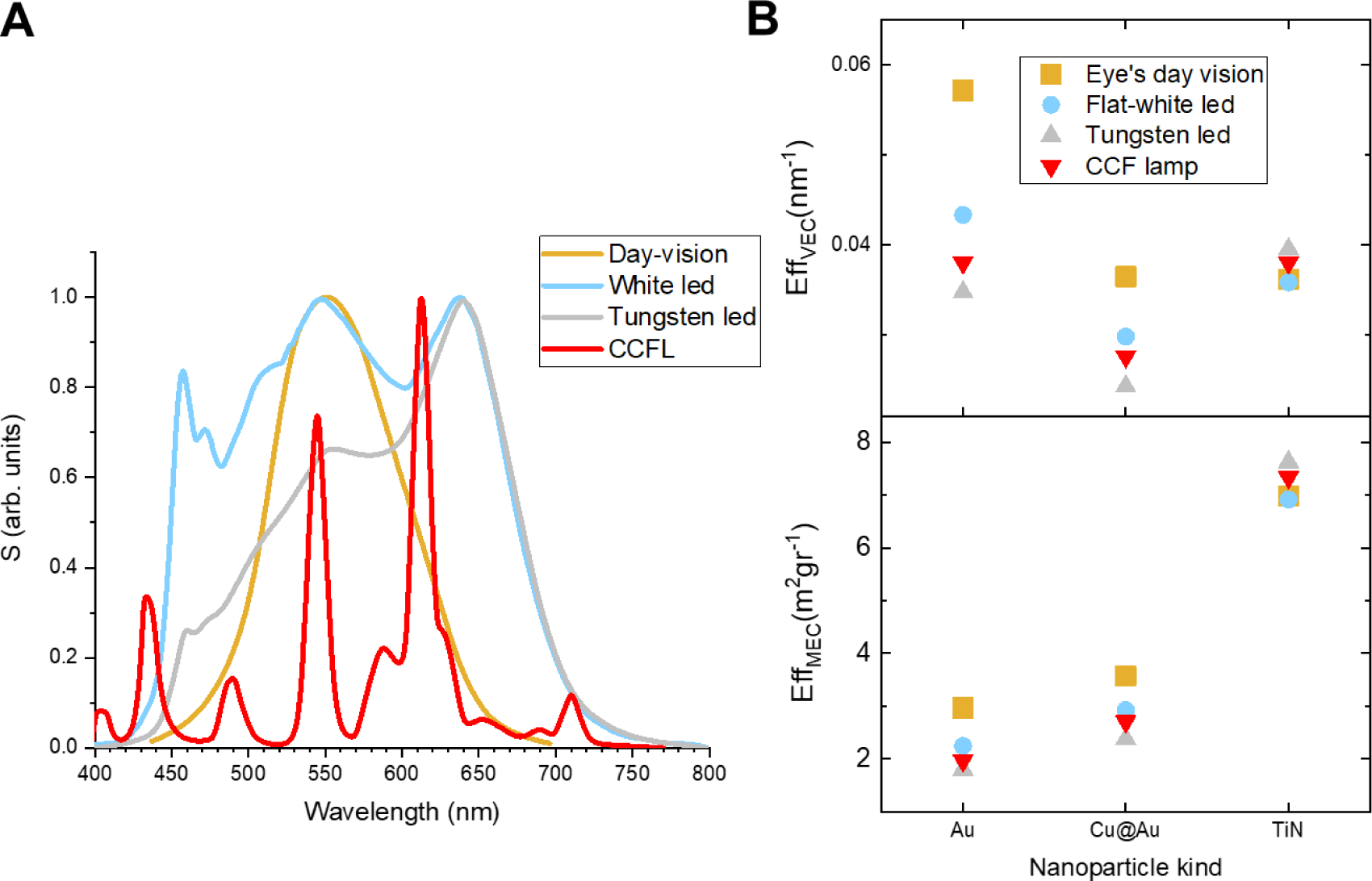
(A) Four different assessments *S*(*λ*) spectra: the day-vision sensitivity of the human eye (photopic vision), a typical tungsten LED, a typical white LED, and cold cathode fluorescent lamp (CCFL). (B) The VEC and the MEC efficiencies for each kind of NP as shown, for the four spectral sources from panel A

#### II.4 Machine learning approach

Our approach incorporates feature extraction from the Red-Green-Blue (RGB) color channels of both the test and control lines on the strips. Colors are represented through combinations of the red, green, and blue primary colors, depicted in Figure S3A.

In the first machine learning experiment, a group-wise k-nearest neighbor (k-NN) model is employed to assess the discriminative capability of the NPs. Within each NP group (TiN, Cu@Au, Au), a pair of strips with the same concentration is randomly assigned for either training or testing, resulting in six strips for each set and a total of 64 rounds of experiments in each NP group. We use k = 1 in the k-NN algorithm, as each class consists of only one sample (strip).

In the second machine learning experiment, we employed contrastive learning to extract features. Contrastive learning uses negative sampling and contrastive loss functions to effectively distinguish between different classes or categories of data. In our experiment, we grouped the strips based on their FITC concentrations. Our contrastive learning model employs a Siamese network ^5^ consisting of two identical subnetworks, each with a simple architecture that includes a single fully connected layer. This Siamese network processes a pair of strips from the training set, generating two corresponding vectors, as shown in Fig. S3B.

**Figure S3.**
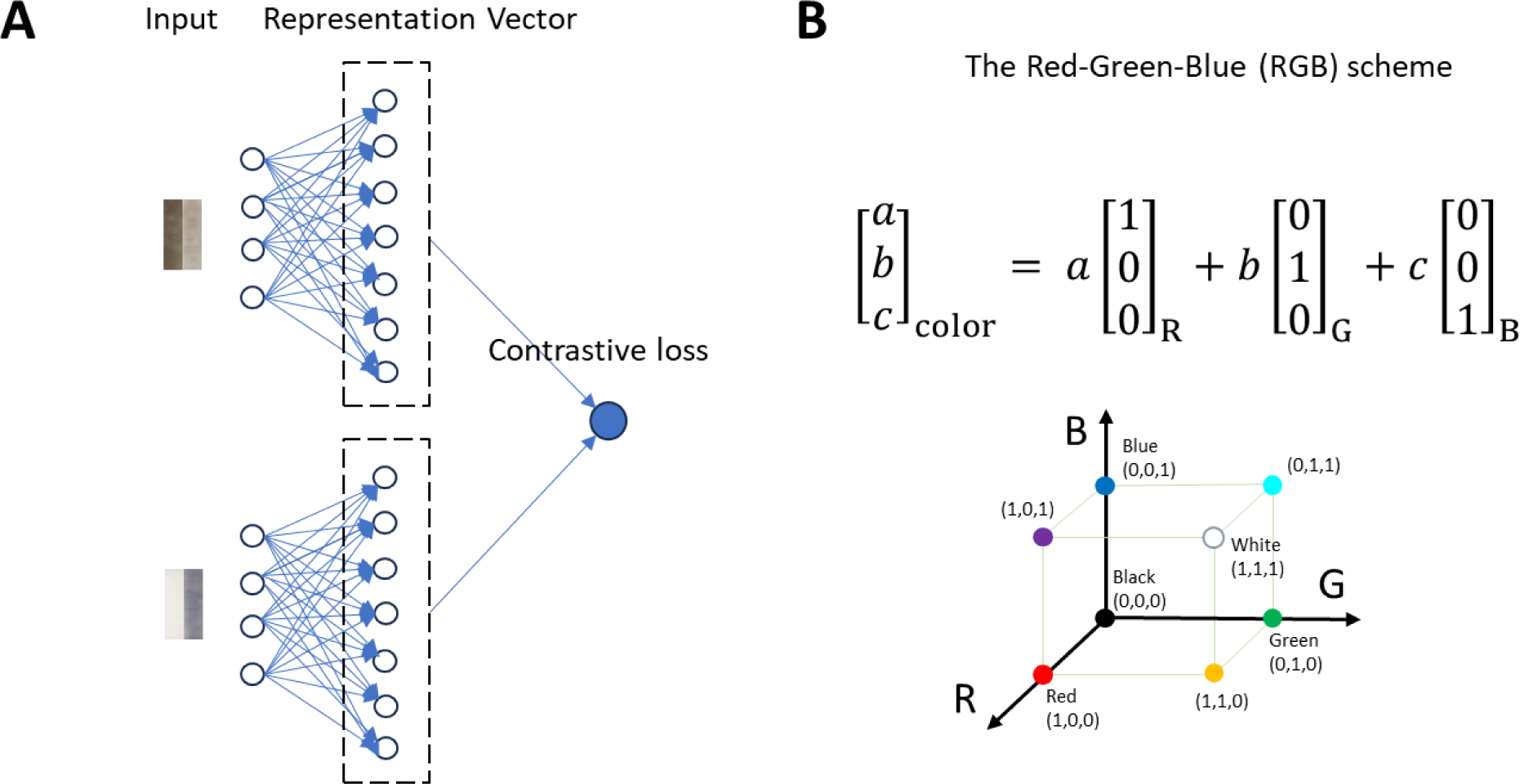
Data, models, and results of our machine learning experiments. (A) Siamese network-based contrastive learning method. (B) Red-Green-Blue (RGB) color channels combination scheme used in this work for feature extraction.

#### III. NPs and transfer to water characterization

**Figure S4.**
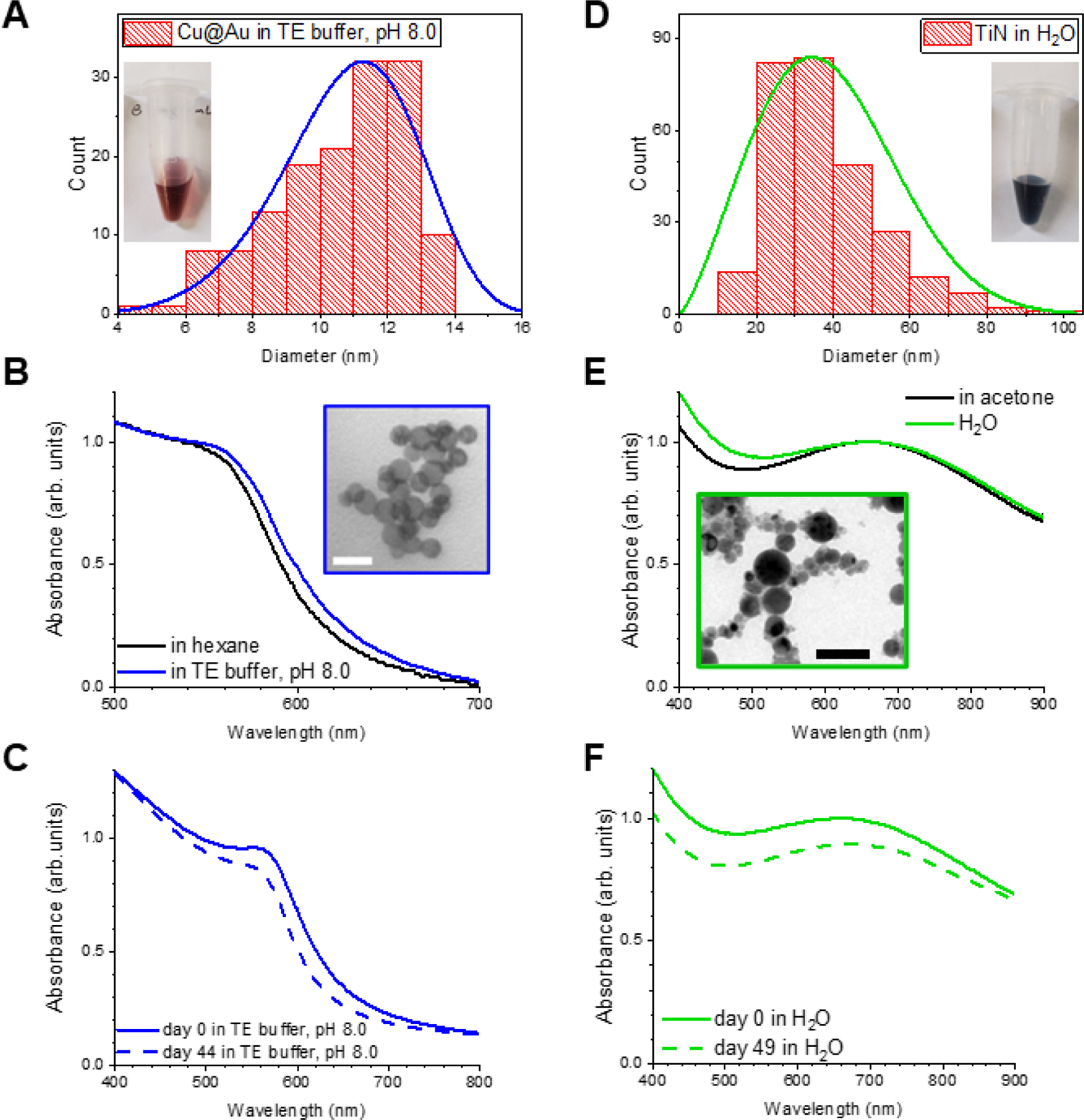
Characteristics of Cu@Au and TiN NPs after their transfer to bio-compatible aqueous media. (A) Plot showing the TEM size distribution of the Cu@Au NPs after their transfer to an aqueous medium. The inset shows a visual of the post-transfer solution. (B) Plasmon of Cu@Au NPs in hexane and aqueous medium. The inset shows a representative TEM image displaying that the NPs show minimal aggregation after being transferred to TE buffer (Bar 20 nm). (C) Conservation of the plasmon through time of the Cu@Au NPs dispersed in TE buffer. (D) Plot showing the TEM size distribution of the TiN NPs after their transfer to water. The inset shows a visual of the post-transfer solution. (E) Comparison of TiN’s plasmon in acetone versus water. The electron microscopy inset confirms that the NPs do not aggregate after acetone evaporation (Bar 100 nm). (F) Conservation of plasmon through time of the TiN NPs.

#### IV. Visible and Near-IR plasmonic characterization for the FITC study

An important characteristic of these NPs is their plasmon, which should be preserved if the NPs want to be used in other approaches, as for example in the near-IR window above the day vision spectrum shown in Figure 1C. To confirm the plasmonic characteristics of the Cu@Au and TiN NPs are maintained after functionalization and LFA, we used the Odyssey XF Imager. This equipment has 3 channels, 685 nm, 785 nm, and 600 nm. Images of the LFAs were taken using each channel. If the NPs preserve their plasmonic characteristics, they will maintain the ability to absorb in the visible and near-IR ranges, and a dark line should be observed. As shown in Figure S5, the images of each channel show a dark line which correlates with the location and intensity of the control and testing line of the naked eye LFAs (showed in the main text on Figure 2C). Moreover, the intensity of the bands observed for each NPs in each channel, correlates with their plasmonic peak (Figure 2B), which confirms that the plasmonic characteristics of these NPs even after functionalization and attaching to the LFAs, is preserved.

**Figure S5.**
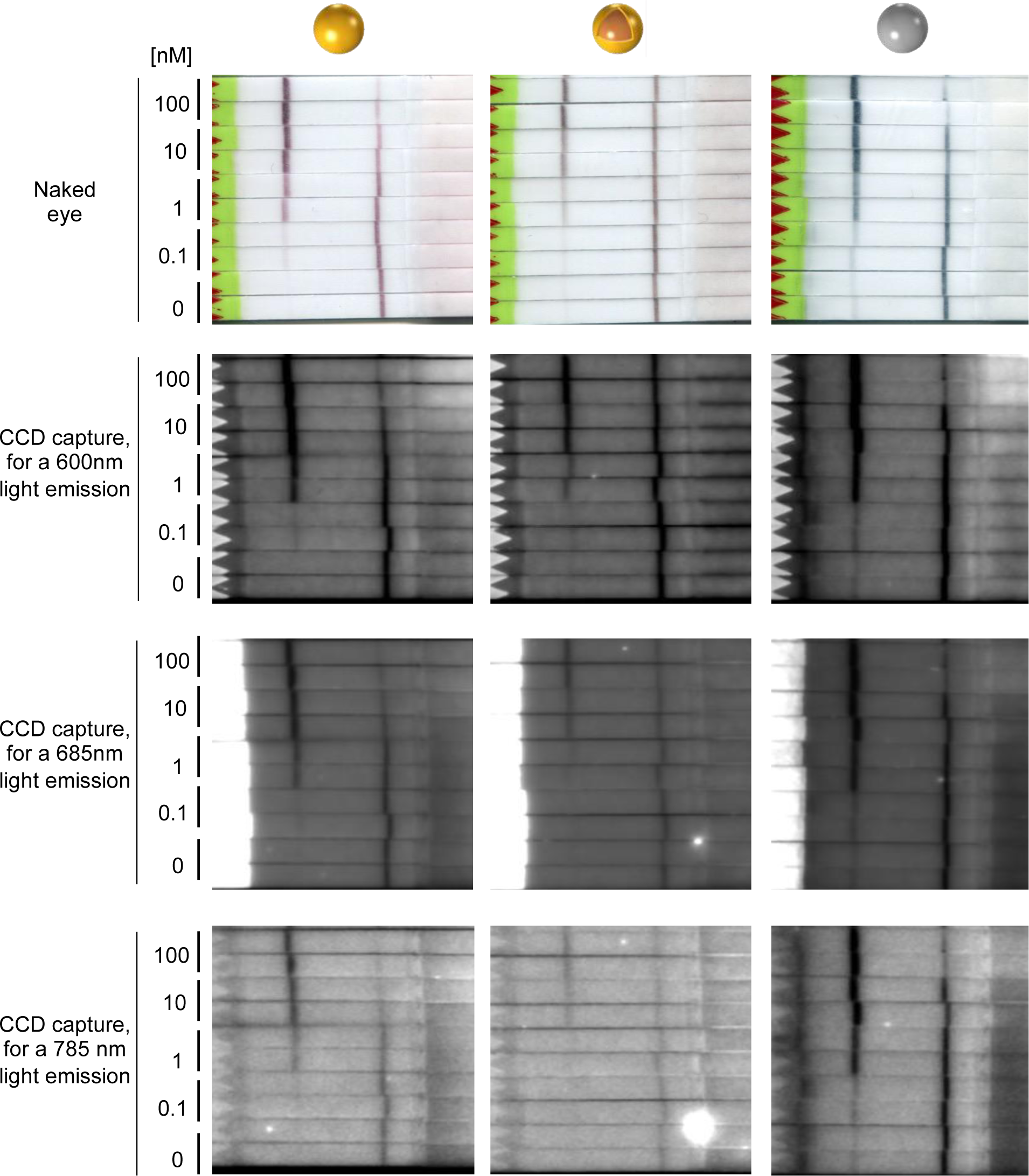
NPs preserve their plasmonic characteristics after functionalization and LFA. The LFAs on Figure 3C were imaged using the Odyssey XF Imager. The LFAs images were taken using each channel (600 nm, 685 nm, and 785 nm) to analyze if the NPs preserved their plasmonic characteristics. As seen in these images, a dark line is observed in each LFAs, which correlated with the images obtained at naked eye. In addition, the images show that each NPs absorb different on each channel, which correlates with their absorption peak characterized in Figure 2B.

#### V. NPs post-functionalization

TEM imaging was developed to confirm that after functionalization the NPs are maintaining in suspension. We use both Cu@Au with the antibody anti-cTnT and TiNs with DNA as examples. The stability of the NPs in an aqueous medium after functionalization is of great importance for the future applicability of these NPs.

Figure S6A was obtained as indicated in the main text, in the materials and methods section. In Figure S6B, the TiN NPs were functionalized using a different approach, to demonstrate the flexibility of conjugation these NPs have. TiN NPs shown in Figure S6B were functionalized with DNA, composed of a thiol group at the 5’ end of 19 consecutive thymine bases, ordered from biomers, that will react with the titanium forming a titanium sulfide bridge between the TiN NPs and the DNA. The method for functionalization was instant dehydration in butanol ^6^. Figure S6B shows that even with this different protocol of functionalization, the TiN NPs do not aggregate. The images were taken using the TEM JEM-1011 transmission electron microscope from JEOL.

**Figure S6.**
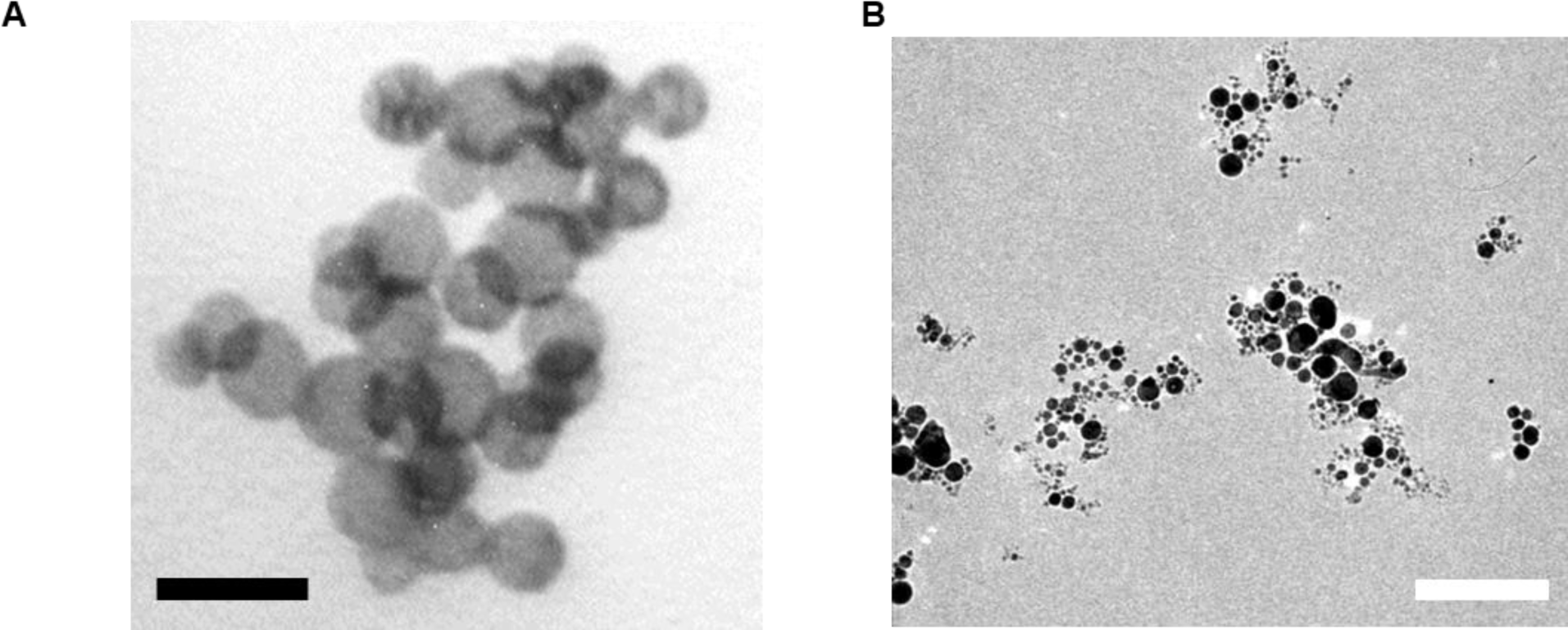
TEM images of NPs after functionalization. A) TEM image of the Cu@Au NPs conjugated with the antibody anti-cTnT (bar 20 nm). B) TEM image of the TiN NPs functionalized with DNA (bar 200 nm). This image demonstrates that different approaches can be used for successfully conjugating the NPs. In this case, DNA composed of 19 thymine based was utilized.

#### VI. Plasmon shift for anti-cTnT functionalization

**Figure S7.**
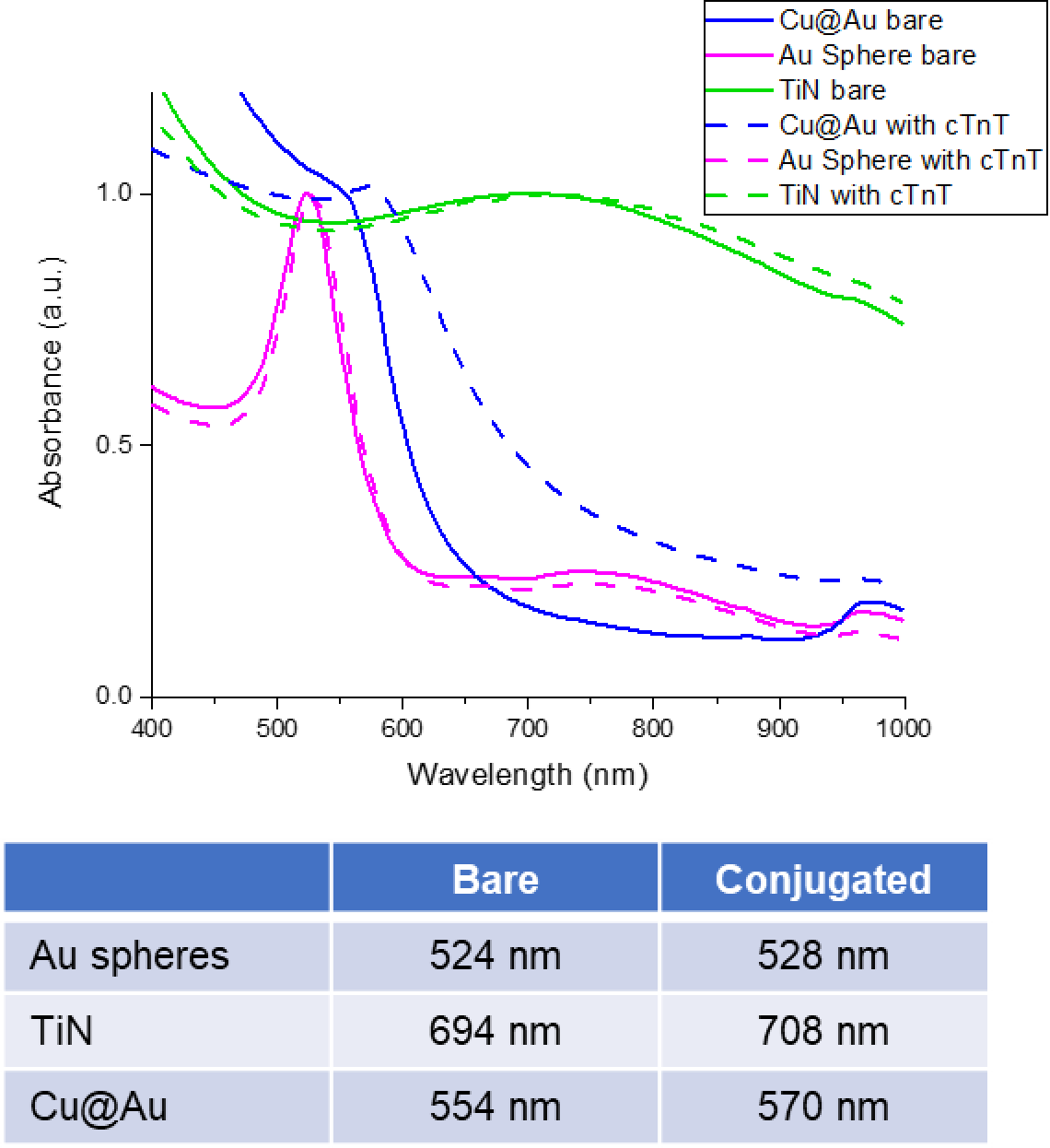
Normalized absorbance of the Cu@Au and TiN NPs pre and post anti-cTnT conjugation. Here, we can see the plot and a table comparing the plasmons for each NPs before (solid lines) and after (dashed lines) conjugation with the anti-cTnT antibody. The results support the successful functionalization of each NPs with the antibody.

